# Robust Inference of Phage-Host Infection Dynamics from Sparse Single-Cell Transcriptomics Profiles in *Pseudomonas aeruginosa*

**DOI:** 10.64898/2026.09.24.752076

**Authors:** Hans Wilms, James Gurney

## Abstract

Phage therapy is providing a benefit to a limited number of people. To date those who have received phage therapy typically have done so via an IND expanded access mechanism, colloquially referred to as compassionate release, new clinical trials are underway, and the advent of phage therapy seems to be at hand. However, a recurring question remains, which phages to use for which bacteria. By understanding how a phage interacts with its bacterial host we will move closer to being able to provide an answer to this pressing question. By using bacterial single cell RNA sequencing (scRNAseq) during phage infection we can look at the most fundamental interaction between the phage and its host. We have observed phage gene expression consistent with well described expression profiles seen in bulk expression data. Phage genes in our scRNAseq data followed the early, mid, and late gene expression profile. Using a machine learning algorithm (Support Vector Machine) we were able to distinguish time of infection from phage genes. We also observed that phage infection is likely not random. When infected with two genetically distinct phages the likelihood of finding both inside a single cell was not consistent with Poisson distribution.

## Introduction

Phage therapy relies on providing a bacterio(phage) that can kill a pathogen. Currently, potential phages are selected based on plate assays used to determine host range and efficiency of plating ^1,2^. These methods provide an accurate but coarse grain estimation of infectivity. However, these assays miss information that could impact treatment. Phage resistance can appear to be a stochastic process, ranging from standing variation of phage receptor mutants in a population, cellular heterogeneity in genes responsible for phage resistance mechanisms (CRISPR/cbass, etc) or even simple physiological states such as growth rate ^3–5^. By examining single cell level data of phage infected bacteria, a better estimation of infectivity can be achieved.

*Pseudomonas aeruginosa* (PA), an opportunistic pathogen that causes severe infections in immunocompromised individuals, is particularly problematic in people with Cystic Fibrosis (pwCF), where chronic infections lead to high mortality ^6^. Due to the bacterium’s innate and acquired resistance to antibiotics ^7,8^ PA infections are difficult to clear, with most pwCF becoming chronically infected by adulthood ^9^. PA isolates from a single pwCF typically show high levels of variance, a single clonal lineage colonizes but diversifies during the decades-long infections ^10^. In environments with antibiotics it is thus inevitable that increased antibiotic resistance will be selected ^12,13^. Treating these chronic antibiotic-resistant infections therefore requires alternative therapies. Phage therapy has gained interest as just one of these alternatives for its abilities in combating antibiotic-resistant bacterial infections ^14,15^. Antibiotic resistance mechanisms are typically distinct from ways in which bacteria can become resistant to phages. However, as with selection for antibiotics, resistance to phages is also likely increased within a population with high diversity, allowing for selection of phage resistant bacterial isolates. Current population level plate-based assays, such as cross streaking, are inadequate to study and identify exactly which members of a bacterial population will be killed by a phage or cocktails of phages. Understanding the infection dynamics at the single cell level would provide far greater resolution for designing and testing phage therapeutics as well as determining the ways in which phage therapy might fail.

Understanding the intricacies of phage-host interactions, especially at the single-cell level, remains a challenge. While bulk RNA-sequencing has been used to analyze the transcriptomic changes in bacterial populations during phage infection ^16^, this method fails to capture the cell-to-cell variability that could provide insight into the stages of phage infection, bacterial defense mechanisms, and the development of resistance. Single-cell-sequencing technologies offer the potential to uncover the diversity in infection states among individual bacterial cells. Single-cell RNA sequencing (scRNA-seq) has been instrumental in uncovering cellular heterogeneity in eukaryotic organisms but has seen less attention in prokaryotes ^17,18^. Small cell size and tough cell walls combined with non-polyadenylated, minimal mRNA content, and an abundance of ribosomal RNA has made this technique particularly difficult in bacteria ^17,19^.

Regardless of the challenges, in recent years, multiple groups have successfully adapted single-cell-sequencing techniques for use in bacterial systems. M3-Seq and BacDrop employ droplet-based technology to generate sequencing libraries, whereas MicroSPLiT and PETRI-seq utilize in situ combinatorial indexing. Although droplet-based methods are faster, they rely on specialized equipment and proprietary commercial kits. In contrast, in situ combinatorial indexing approaches are more accessible, requiring only standard laboratory equipment such as a centrifuge and thermocycler, without the need for commercial kits ^20–23^.

PETRI-seq, a prokaryotic single-cell RNA sequencing technique, allows for the study of bacterial cells at the individual level, offering high resolution in understanding host-pathogen interactions ^21^. PETRI-seq uses *in situ* combinatorial indexing with high-throughput sequencing, enabling simultaneous profiling of thousands of individual cells. Unlike methods like ProBac-seq, PETRI-seq uses a non-targeted approach so that prior knowledge of bacterial or phage genetic content is not necessary ^24^. This method has the potential to revolutionize the study of phage biology by revealing how bacterial cells within a population respond heterogeneously to phage infections.

In this study, we applied a modified version of PETRI-seq to investigate the infection dynamics of Luz19 and LKD16 phages in PA cells. Although single-cell transcript capture in *Pseudomonas* was sparse, we were still able to detect structured phage infection signals across multiple time points, classify infection states using machine learning, and identify genes associated with infection progression. In mixed infections, Luz19 and LKD16 were detected together in the same cells more often than expected, suggesting that single-cell approaches can reveal phage–host and phage–phage dynamics that may be missed by bulk or plate-based assays.

## METHODS

### Bacterial and phage strains and growth conditions

*P. aeruginosa* PAO1 was grown in 50ml conical tubes in LB medium at 37°C and shaken at 165r.p.m. until log phase. Cultures were infected with Luz19 phage at an MOI of 5 and further incubated for 1hr. Timepoints were collected at -1min pre-infection and 10, 20, 30, and 40min post-infection.

For coinfection experiments, PAO1 was infected with Luz19 phage at an MOI of 0.57 and LKD16 phage at an MOI of 0.35. The infection continued at room temperature and timepoints were collected at 5, 10, 15, and 20min post-infection.

### PETRI-seq

Cell and library preparation for PETRI-seq was performed according to the original PETRI-seq method^21^ with some modifications to improve recovery of PAO1 cells (faster centrifugation and addition of Tween20) and reduce overabundance of rRNA reads (rRNA depletion). All centrifugation was performed at 10,000g for 10 minutes at 4°C (Sorvall Legend Micro 21R centrifuge) unless otherwise noted.

### Barcode plate preparation and annealing

Round 1, 2, and 3 barcode oligos, linker oligos, and blocking oligos were ordered and prepared according to the methods originally described in the original PETRI-seq method^21^.

### Cell fixation and permeabilization

For each timepoint, 1ml of culture was harvested by centrifugation for 5 minutes. After the supernatant was removed, cells were resuspended in 15mL conical tubes using 3mL of ice-cold 4% formaldehyde (Sigma-Aldrich, catalog no. F8775) in 1x PBS (Thermo Fisher Scientific, catalog no. 10010023) and rocked for ∼16 hours at 4°C (Ohaus RockingShaker).

Following fixation, cells were divided centrifuged in microcentrifuge tubes and the supernatant was discarded. The cell pellets were washed by resuspension in 1mL of PBS-RI-Tween (1x PBS supplemented with 0.01 U/µL SUPERase In RNase Inhibitor, Thermo Fisher Scientific, catalog no. AM2696 and 0.01% Tween20, Sigma-Aldrich, catalog no. P9416) and centrifuged again and the resulting pellets were resuspended and pooled in 300 µL of ice-cold PBS-RI-Tween.

After centrifugation, the cells were resuspended in 150µL of PBS-RI-Tween, and 150µL of ice-cold ethanol was added. The cells were washed twice with 300µL of ice-cold PBS-RI and counted using a Petroff-bacteria counter. 4×10^7^ cells were transferred to a PCR tube and suspended in 100 µL of lysozyme semi-permeabilization buffer (100mM Tris pH 8, 50mM EDTA, 2.5mg/mL lysozyme, and 0.25U/µL SUPERase In RNase Inhibitor). The permeabilization was carried out at 37°C for 15 minutes in a pre-warmed thermocycler. After permeabilization, cells were transferred to 1ml PBS-RI-Tween, washed twice with 300 µL PBS-RI.

### *In situ* ribosomal depletion

The *in situ* ribosomal depletion was carried out similar to Ma *et al*^*22*^ using the NEBNext rRNA Depletion Kit (Bacteria) (NEB, catalog no. E7850L). Cells were resuspended in 11µL water and transferred to a PCR tube. 2µL of rRNA depletion solution and 2µL of probe hybridization buffer was added, and solution was incubated in the thermocycler with the lid set to 55°C. Cells were incubated at 50°C for 2min, then ramped down ∼0.1°C/sec to 22°C, then a 5min hold at 22°C. As soon as the 5min hold was finished, 2µL RNase H and 2µL reaction buffer and 1µL water was added, and cells were further incubated at 50°C for 30min. After incubation, 180µL ice-cold PBS-RI-Tween was added to the solution and centrifuged at 12,000G for 15min at 4°C. DNase digestion was performed by resuspending pellet in 16.4µL nuclease free H2O, 0.65µL 1:50 diluted SUPERase In RNase Inhibitor, 1µL 10X DNase buffer, and 1µL DNase, then incubating at 25C for 30min. DNase was inactivated by addition of 1µL 50mM EDTA and incubating at 50°C for 10min.

180uL of PBS-RI-Tween was added and cells were transferred to a microcentrifuge tube. Cells were washed twice in 100uL PBS-RI-Tween and resuspended in 20uL PBS-RI-Tween before continuing to Split-Pool Barcoding.

### Split-Pool Barcoding

To multiplex infection timepoints within a single PETRI-seq experiment, cells from each timepoint were assigned to a distinct subset of wells during the first round of combinatorial barcoding. For example, cells from timepoint 1 were distributed across rows A–B of the first barcoding plate, timepoint 2 across rows C–D, timepoint 3 across rows E–F, and timepoint 4 across rows G–H. Thus, each timepoint occupied 24 wells of the 96-well first-round barcoding plate.

This design allowed the first-round barcode to serve both as part of the combinatorial cell barcode and as a timepoint identifier. Because each timepoint was assigned a non-overlapping set of first-round barcode wells, cells from different timepoints could be distinguished downstream based on their first-round barcode identity while being processed together in subsequent PETRI-seq steps.

Cells were split into a 96-well PCR plate containing Round 1 barcoded reverse transcription (RT) oligonucleotides. An RT mix (240 µL 5x Maxima H Minus RT buffer, 24 µL 10 mM dNTPs, 12 µL SUPERase In RNase Inhibitor, 24 µL Maxima H Minus Reverse Transcriptase, and 640 µL RNase-free water) was prepared and 8 µL of the mix was added to each well containing cells and 2 µL of oligos. The RT reaction was performed with the following thermocycling conditions: 50°C for 10 minutes, 8°C for 12 seconds, 15°C for 45 seconds, 20°C for 45 seconds, 30°C for 30 seconds, 42°C for 6 minutes, and 50°C for 16 minutes, followed by a 4°C hold.

Following the RT step, cells were pooled, and Tween-20 was added to a final concentration of 0.01%, followed by centrifugation at 20 minutes. Cells were resuspended in 600 µL of T4 ligase buffer supplemented with 0.1 U/µL SUPERase In RNase Inhibitor.

For Round 2 barcode ligation, following centrifugation, the supernatant was carefully removed, and the cells were resuspended in 600 µL of 1x T4 ligase buffer (Thermo, catalog no. EL0011), supplemented with 0.1 U/µL SUPERase In RNase Inhibitor.

A ligation master mix was then prepared by combining 7.5 µL of RNase-free water, 37.5 µL of 10x T4 ligase buffer, 16.7 µL of SUPERase In RNase Inhibitor (20 U/µL), 5.6 µL of BSA (New England Biolabs, catalog no. B9001S), 27.9 µL of T4 ligase (Thermo, catalog no. EL0011), and 600 µL of the resuspended cells. 5.76 µL ligation master mix was aliquoted into each well of a 96-well plate containing 2.24 µL of pre-aliquoted Round 2 annealed oligonucleotides.

The ligation reaction was incubated at 37°C for 30 minutes. Following the ligation step, a Round 2 blocking mix was prepared and 2 µL of this blocking mix was added to each well. The Round 2 blocking mix was composed of 37.5 µL of 400 µM SB84 blocking oligonucleotide, 37.5 µL of 400 µM SB85 blocking oligonucleotide, 25 µL of 10x T4 ligase buffer, and 150 µL of RNase-free water. The blocking reaction was incubated at 37°C for an additional 30 minutes. After this incubation, the reactions were pooled into a single tube.

For the Round 3 barcode ligation, 46 µL of 10x T4 ligase buffer, 12.65µL of T4 ligase, and 115µL of RNase-free water was added to the pooled cells. 8.51µL of this ligation mix was added to each well of a 96-well plate which already contained 3.49 µL of pre-aliquoted Round 3 annealed oligonucleotides. The ligation reaction was incubated at 37°C for 30 minutes.

After the ligation, 10µL of a Round 3 blocking mix was added to each well. The Round 3 blocking mix was composed of 72µL of 400 µM SB81 blocking oligonucleotide, 72 µL of 400 µM SB82 blocking oligonucleotide, 120 µL of 10x T4 ligase buffer, 336 µL of RNase-free water, and 600 µL of 0.5 M EDTA. After adding the blocking mix, the cells were pooled, tween-20 was added to a final concentration of 0.01%, and the cells were centrifuged. The supernatant was removed, and the pellets were resuspended in 500 µL of TEL-RI buffer (100 mM Tris pH 8.0, 50 mM EDTA, and 0.1 U/µL SUPERase In RNase Inhibitor) with 0.01% Tween-20.

The cells were counted using a semen test hemocytometer (Hausser Scientific, catalog no. 3900). Approximately 10,000 cells were transferred into individual microcentrifuge tubes for lysis. The volume was adjusted to 25 µL using TEL-RI buffer and lysed by adding 25 µL of 2x Lysis-T buffer (50 mM EDTA, 400 mM NaCl, 1% Triton X-100) and 5 µL of proteinase K (20 mg/mL, Thermo Fisher Scientific, catalog no. AM2548) to each tube. The cells were lysed at 55°C for 1 hour with occasional agitation. After lysis, the samples were stored at -80°C for future use.

### Library Preparation and Sequencing

Libraries were prepared according to the methods originally described in the original PETRI-seq method^21^. Sequencing was performed across three runs, designated JRG06, JRG07, and JRG09. PETRI-seq libraries concentrations were quantified using a DeNovix DS-11 FX+ fluorometer with the High Sensitivity assay kit. Libraries were pooled in equimolar proportions, and the resulting pools were diluted and denatured according to the standard Illumina sequencing protocol. The JRG05 and JRG06 pools were sequenced on an Illumina MiniSeq using the MiniSeq Mid Output Kit with paired-end 100 bp (PE100) reads. The JRG07 and JRG09 pools were sequenced on an Illumina NextSeq 500 using the Mid Output and High Output kits, respectively, with paired-end 75 bp (PE75) reads to achieve increased sequencing depth.

### PETRI-seq pipeline and analysis

Raw paired-end sequencing reads generated from PETRI-seq libraries were processed using a custom computational pipeline based on the original PETRI-seq workflow^21^. Read 1 was used to extract the in situ barcodes (BC1–BC3) and unique molecular identifier (UMI), while Read 2 contained the cDNA sequence. Reads were filtered for quality and trimmed to remove adapter sequences using cutadapt. UMIs were parsed from Read 1 using umi_tools extract and cutadapt was used to demultiplex barcodes 3, 2, and 1 into separate FASTQ files in a custom Python script. For timepoint samples split on BC1 plates, cells were computationally assigned to infection timepoints using the first-round barcode index from the final PETRI-seq cell barcode.

A cumulative frequency table was built using python script and a number of barcodes were selected based on a knee plot of barcodes vs cumulative reads so that only barcodes with a high number of reads were kept.

Read 2 sequences were aligned to the Pseudomonas aeruginosa PAO1 reference genome (RefSeq NC_002516) using BWA. Aligned reads were sorted and indexed using SAMtools. Gene-level assignment was performed using featureCounts, excluding multi-mapping reads. UMI deduplication and cell-by-gene count matrix generation were conducted using UMI-tools v0.5.5, ensuring that PCR duplicates were collapsed based on identical UMIs mapped to the same cell.

Following matrix construction, low-quality cells were filtered prior to downstream analysis (only cells expressing more than 5 genes were kept). Dimensionality reduction, clustering, and visualization were performed using **Scanpy**, including normalization, log transformation, and principal component analysis (PCA), followed by Uniform Manifold Approximation and Projection (UMAP) for embedding. Cell clusters were identified using the Leiden algorithm, and differentially expressed genes were computed using Scanpy’s rank_genes_groups function.

### Phage adsorption

Overnight PAO1 cultures were grown to log phase, counted on a Petroff-Hausser chamber, and infected at the indicated MOI in 5ml of LB media.

## Results

PETRI-seq was performed on 4 different samples using 6000-24,000 cells, with 16-50 million reads obtained per sample (Table Si1). The transcript capture per cell was poor in *Pseudomonas*, with the vast majority of cells harboring less than 12 UMIs, irrespective if rRNA depletion was carried on that sample or not (Fig 1A). Figure 1 shows general information on the reads profile. Total reads ranged from 16-52 million reads per experiment, or 796-5530 reads per cell. Given the high duplicate read rates (Fig 1B) and stifled saturation curves (Fig 1D), it does not seem like more sequencing would have been beneficial. The distribution of reads per UMI indicates over-cycling during the library preparation (Fig 1C), which was necessary to reach the minimal amount of input for sequencing.

**Figure 1.**
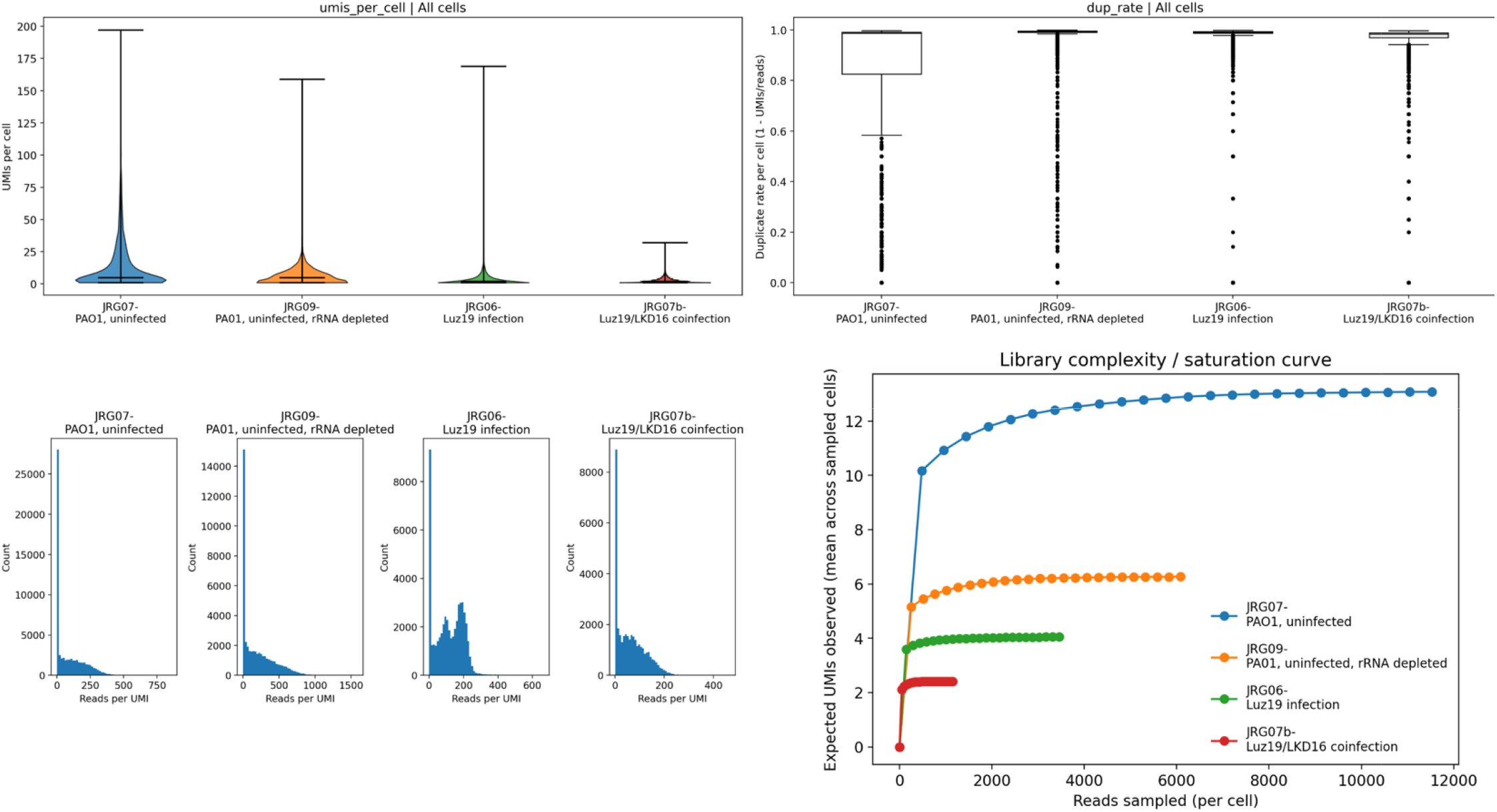
Splitseq of phage infected bacteria produced low yield mRNA read. A. Violin plots of UMI/cell for each experiment. The average UMI count was 12. B. Boxplots showing the distribution of per-cell duplicate rates. Higher values indicate a greater proportion of sequencing duplicates relative to unique UMIs. C. Histograms showing the distribution of read per UMI. D. For each dataset, library complexity is summarized as the expected number of distinct UMIs observed as a function of reads sampled per cell.

While transcript capture was poor, Monte Carlo simulation of gene co-occurrence showed that cell expression was not random (Fig 2A). Infection of PAO1 with Luz19 caused a predictable pattern of infection, with early Luz19 genes primarily being expressed the from the ‘lefthand’ side of the of the genome at 10min post infection and shifting to the ‘righthand’ side expressing late genes at >30min post infection (Fig 2B).

**Figure 2.**
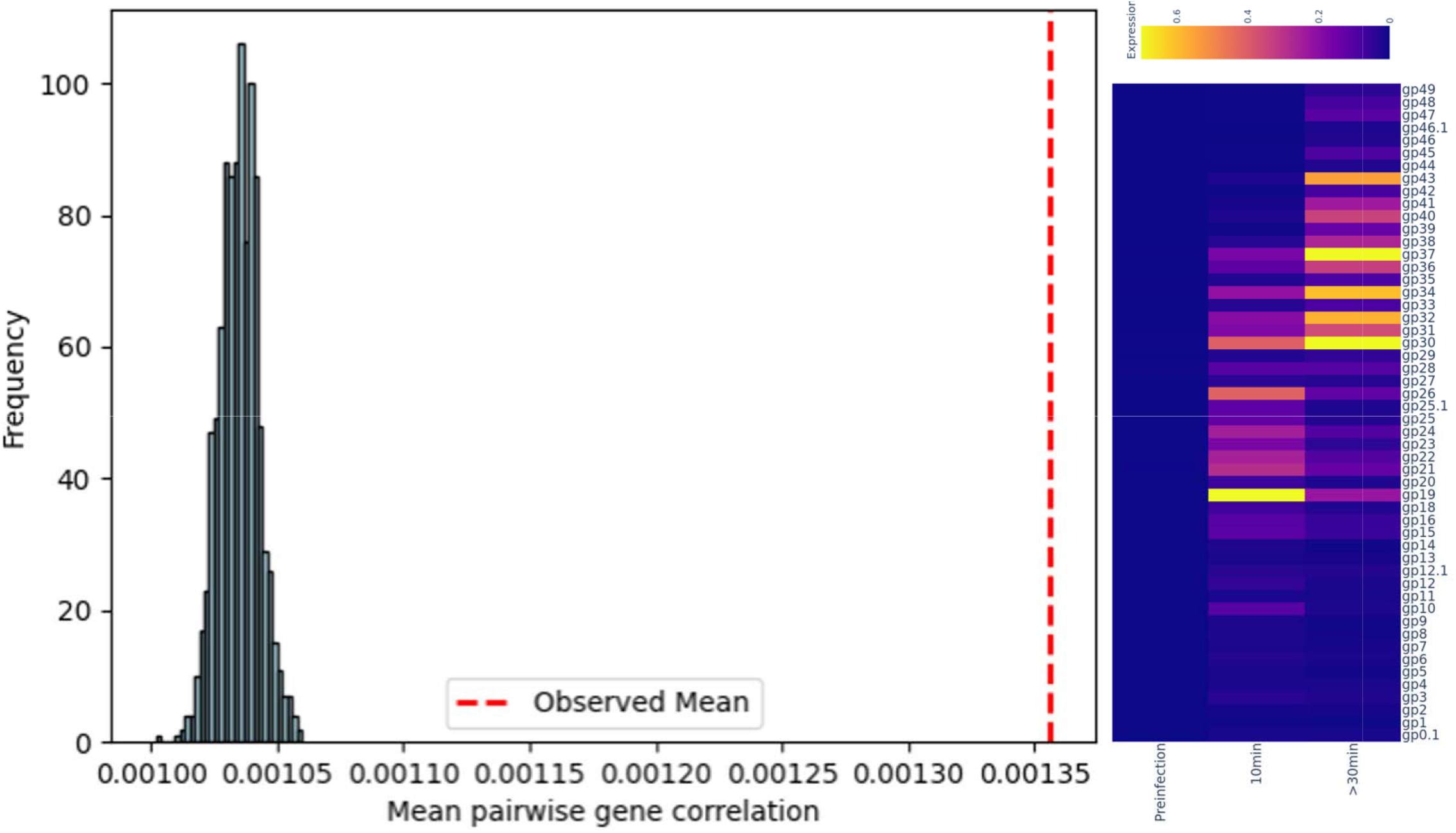
scRNAseq data was non-random and provided expected temporal dynamics. A. Histogram from a Monte Carlo reassortment test showing the distribution of mean absolute pairwise gene–gene correlations expected under a null model where gene expression is randomly shuffled within each cell. Each bar represents the frequency of simulated datasets with a given mean correlation, and the dashed red vertical line indicates the observed mean correlation from the unshuffled data. B. Heatmap showing mean expression of Luz19 genes across cells at each timepoint. Columns represent individual Luz19 genes, ordered by genomic location, and rows represent timepoint groups. Color intensity indicates mean counts per cell.

The Luz19 infected cell expression clustered by timepoint in a UMAP (Fig 3A). More cells from the >30min group were found within the 10min cluster, as these likely represent secondary or later infections that are just getting started. Many cells from the 10min and >30min groups also clustered with the pre-infection cells, and likely represent cells that evaded infection. Due to the magnitude of shift in gene expression during phage infection, SVM analysis was able to classify 10min and >30min cells reasonably well considering the limited dataset (Fig 3B and C). Over 80% of 10min cells and 90% of >30min cells were correctly predicted when trained off 25% of the data. PC1 loadings of the top 15 genes showed Luz19 gene expression of gp19, and gp26 pulling cells to the early group, while expression of later structural genes pushed cells into the >30min group (Fig 3D).

**Figure 3.**
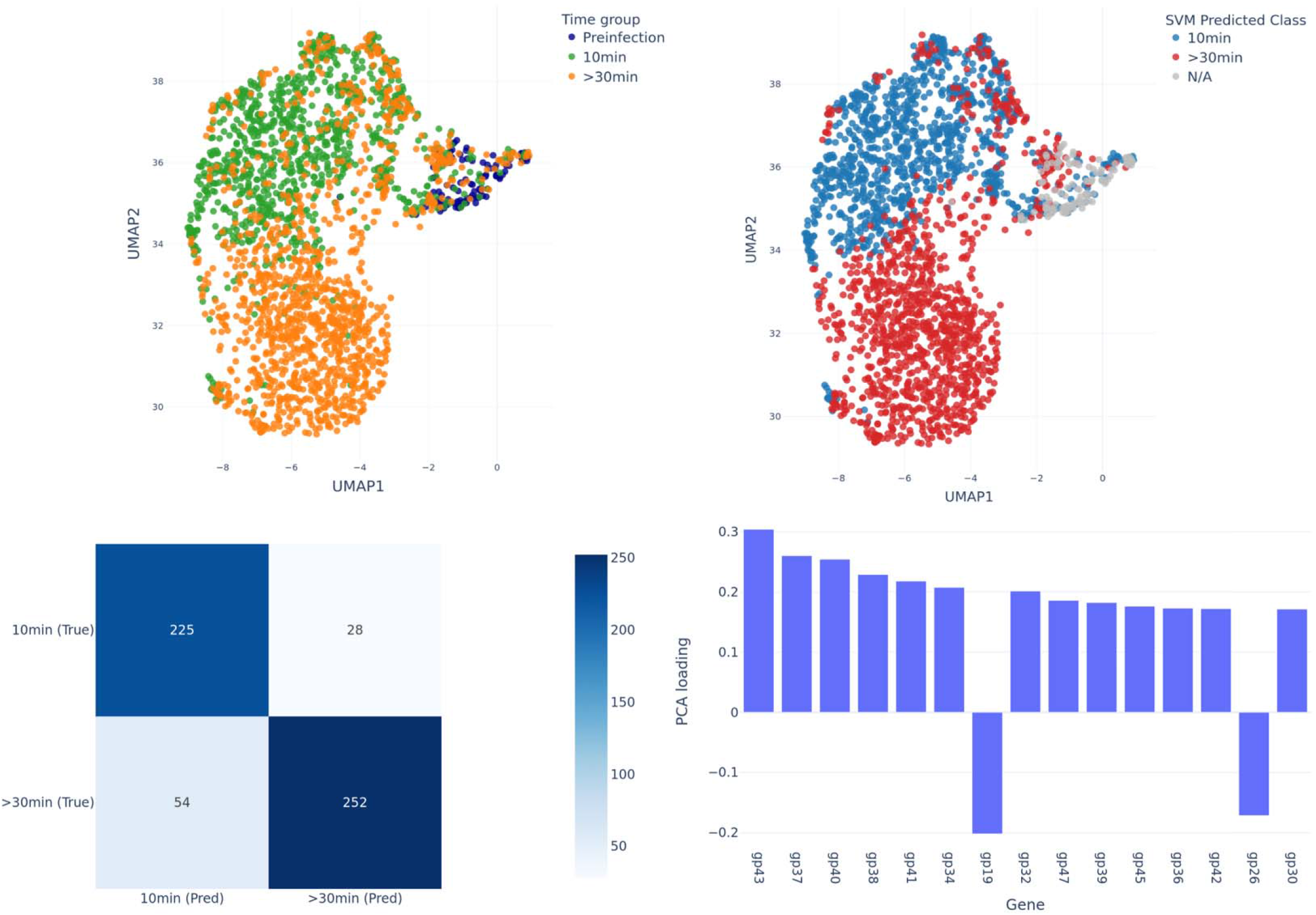
Phage infection status was predictable from mRNA signature. A. UMAP visualization of single cells by infection time group. B. UMAP visualization of single cells colored by SVM-predicted class label. Only 10min and >30min were analyzed by SVM, the preinfection that were not included in SVM-prediction were graphed in grey. C. Confusion matrix summarizing SVM classification performance, showing the number of cells assigned to each predicted class versus their true class label. D. Bar plot of PC1 loadings showing the contribution of individual genes to the first principal component used in the SVM analysis for >30min period.

The co-infection of PAO1 using Luz19 and LKD16 produced unexpected results. Infected cells were lower than expected based on a chi-square analysis during early infection at 5 and 10min post infection. This was simply because substantial viral genes could not yet be detected this early in the infection. However, later during infection when phage genes could be reliably detected (15-20min), co-infected cells were significantly higher than expected based on MOI. Because this infection occurred at room temperature and the nature of temporal gene expression in phages, it is unlikely that these were the result of secondary infections. Strikingly, single cell infections of only Luz19 or LKD16 remained less than expected throughout the infection (Fig 4A). Heat maps at 15 and 20min post infection show co-infected cells seemed to express higher amounts of multiple viral genes than singly infected cells (Fig 4B), but cell counts per gene were too low for proper statistical analysis. This higher expression of viral genes in co-infected cells was not simply due to more raw reads being detected in co-infected cells, as Figure 4D shows. Heatmaps at 5 and 10min post infection were not included due to low number of cells expressing viral genes detected at these timepoints. At 15 min post infection, later phage genes (gp20 and up) seemed to co-express in coinfected cells (Fig 4C). Notably, Luz19 and LKD16 gp26 was high in co-expression with its extra-species partner.

**Figure 4.**
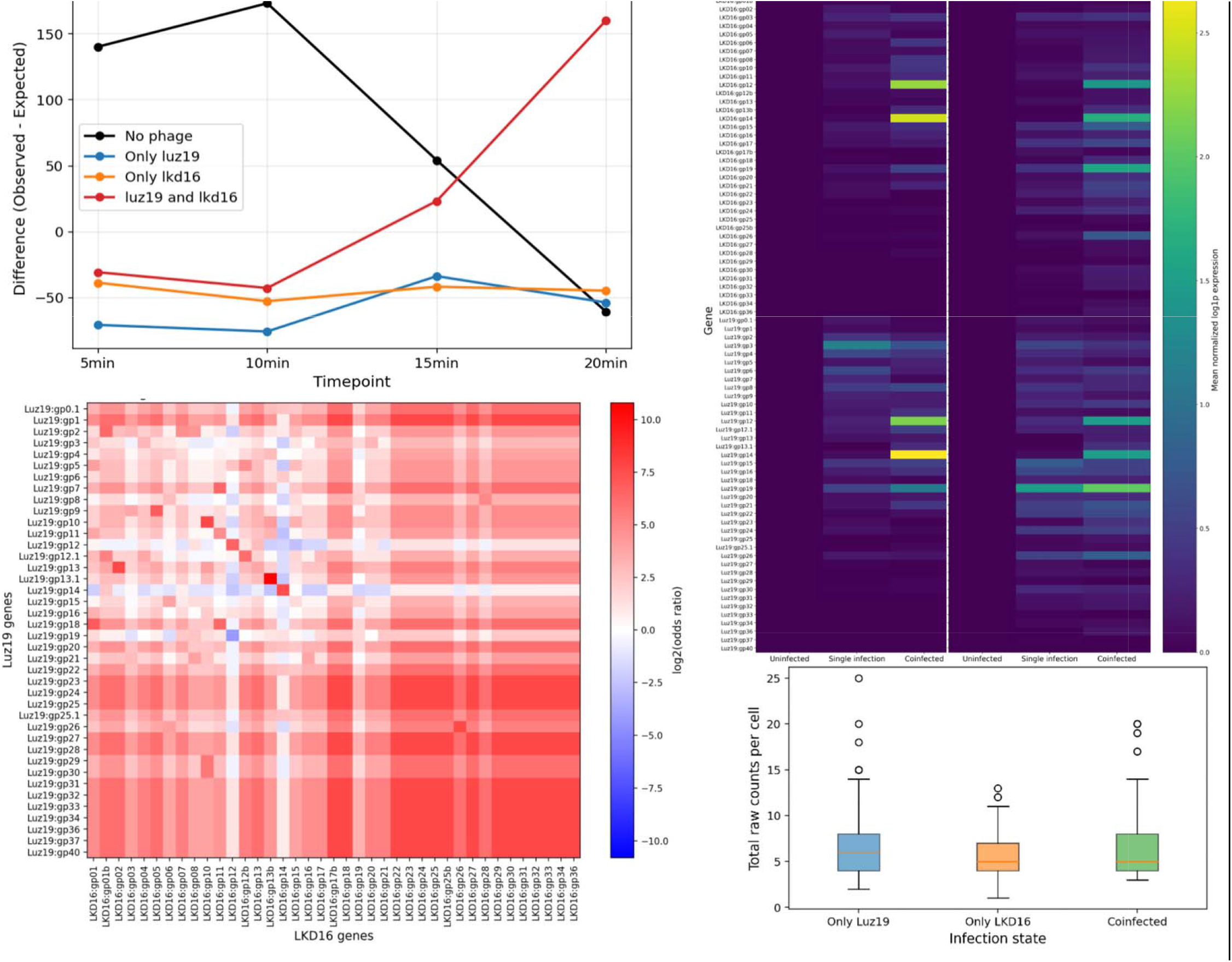
Coinfection of luz19 and LKD16 was detectable with PETRIseq. A. Line plot showing the difference between observed and expected cell counts for each infection state across postinfection timepoints. Positive values indicate more cells than expected under a Poisson MOI model for that population, and negative values indicate fewer cells than expected. B. Heatmaps showing mean gene expression by infection state at 15- and 20-min post infection. Rows correspond to genes and columns to infection states (uninfected, single infection, and coinfected), with color intensity indicating the mean expression level across cells within each group. C. Heatmap depicting the correlation between Luz19 and LKD16 gene expression across single cells. Phage genes are ordered by location on the genome. D. Box plots showing the distribution of total raw transcript counts per cell for each infection state at individual timepoints, with boxes representing the interquartile range, center lines indicating the median, and whiskers denoting the spread of the data.

## Discussion

A major practical goal of this study was to test whether a PETRI-seq–style workflow can recover informative phage infection signals from *Pseudomonas aeruginosa* at single-cell resolution, even when transcript capture is poor. In PAO1, we observed extremely low UMI recovery per cell—most cells contained fewer than ∼12 UMIs regardless of whether *in situ* rRNA depletion was applied. This limited sensitivity was accompanied by high per-cell duplicate rates and library complexity curves that plateaued early, consistent with substantial PCR/sequencing duplication and incomplete molecular diversity. Together, these metrics argue that additional sequencing depth alone would not have rescued information content for these libraries. Instead, future gains will mainly come from improving capture efficiency upstream (permeabilization/retrotranscription/ligation chemistry and amplification strategy). Notably, the median capture reported in the original PETRI-seq work (e.g., hundreds of mRNAs per *E. coli* cell under some conditions) highlights how strain dependent capture can be, and emphasizes that bacteria like PAO1 may require tailored optimization rather than direct transfer of a general protocol ^21^.

Despite low transcript capture, the single-cell phage signal was not random noise. A Monte Carlo shuffle test rejected a null model of randomized within-cell gene expression structure, supporting the presence of real, coordinated transcriptional programs in infected cells. Moreover, when Luz19-infected cell expression was aggregated it exhibited a clear temporal pattern: early infection (10 min) was enriched for transcripts mapping toward one side of the Luz19 genome, with a shift toward later genes in the >30 min group. This is consistent with prior bulk sequencing experiments showing staged, time-resolved transcriptional programs during phage infection ^16^. Importantly, these results support the central “proof-of-principle” conclusion: even sparse bacterial single-cell libraries can still robustly capture phage infection dynamics when phage-driven transcriptional changes are large relative to technical dropouts.

Dimensionality reduction and supervised learning further reinforce that the recovered signal is biologically separated by infection time group, with preinfection cells forming a distinguishable population and infected cells diverging along trajectories consistent with progressive infection states. The presence of >30 min timepoint cells within the 10 min cluster is plausibly explained by infection as a continuous process, as cells harvested at later times can still be at earlier transcriptional stages (e.g., delayed adsorption, slower intracellular progression, or secondary infection). In parallel, many cells from the infected timepoint samples clustered with preinfection cells, indicating cells that evaded infection. Consistent with this interpretation, an SVM classifier trained to separate 10 min vs >30 min infection states achieved success with the limited dataset (∼80% correct for 10 min and ∼90% correct for >30 min). The observed confusion, i.e. late timepoint cells predicted as “early”, may therefore reflect genuine biological overlap, rather than algorithmic error.

The PC loadings provide mechanistic hypotheses about the transcriptional transition captured by the classifier. In the single infection Luz19 dataset, gp19 and gp26 load toward the “early” grouping while later structural genes push cells into the >30 min grouping. Notably, gp26 in phiKMV-like phages encodes single-subunit RNA polymerase (“vRNAP”) responsible for transcription of structural and lysis-associated late genes, and is tied to the shift toward late transcriptional control ^25^. Even with limited bacterial transcript capture, our single-cell data aligns with the idea that gp26 acts as a trigger that separates infection stages, albeit this is only detectable due to the magnitude of effect the phage infection has on the cell’s transcriptional state.

The most unexpected result was from the Luz19 + LKD16 coinfection experiments. At early timepoints (5–10 min), infected cells were lower than expected, which was likely due to detection limits before viral transcripts accumulate to reliably measurable levels. At later timepoints (15–20 min), however, coinfected cells were significantly more frequent than expected under the MOI-based Poisson distribution estimation, while singly infected cells remained below expectation. This pattern argues against a simple model where Luz19 infects permissive cells based on random interaction, and instead suggests either (i) strong host-cell heterogeneity in susceptibility or (ii) phage induced facilitation, where infection by one phage increases the probability of productive infection by the other. The room-temperature infection condition makes rapid secondary rounds of infection less likely, strengthening the case that the enriched coinfection fraction reflects either pre-existing susceptible states or phage induced facilitation rather than serial infection cascades. This is to say, if a bacterial host was permissive to one phage it had a higher likelihood of being permissive to another. Transcript-level patterns within coinfected cells provide additional (but still preliminary) support for interaction rather than simple additivity. At 15–20 min, coinfected cells appeared to express higher amounts of multiple viral genes than singly infected cells. This was not explained simply by higher raw read depth per cell in the coinfected group as assessed by the total raw transcript distributions (Fig 4D). The phage gene– gene correlation heatmap further suggested coordinated expression among sets of later genes between species (Fig 4C). Given that LKD16 and Luz19 are both phiKMV-like phages, a plausible mechanistic hypothesis is that similarities in transcriptional architecture make coinfection synergistic rather than combative. But more robust data is needed for any solid claims.

## References

1. Kutter, E. Phage host range and efficiency of plating. Methods Mol. Biol. 501, 141–149 (2009).

2. Khan Mirzaei, M. & Nilsson, A. S. Isolation of Phages for Phage Therapy: A Comparison of Spot Tests and Efficiency of Plating Analyses for Determination of Host Range and Efficacy. PLOS ONE 10, e0118557 (2015).

3. Chapman-McQuiston, E. & Wu, X. L. Stochastic receptor expression allows sensitive bacteria to evade phage attack. Part I: experiments. Biophys. J. 94, 4525–4536 (2008).

4. Turkington, C. J. R., Morozov, A., Clokie, M. R. J. & Bayliss, C. D. Phage-Resistant Phase-Variant Sub-populations Mediate Herd Immunity Against Bacteriophage Invasion of Bacterial Meta-Populations. Front. Microbiol. 10, 1473 (2019).

5. Hobbs, S. J. & Kranzusch, P. J. Nucleotide Immune Signaling in CBASS, Pycsar, Thoeris, and CRISPR Antiphage Defense. Annu. Rev. Microbiol. 78, 255–276 (2024).

6. Malhotra, S., Hayes, D. & Wozniak, D. J. Cystic Fibrosis and Pseudomonas aeruginosa: the Host-Microbe Interface. Clin. Microbiol. Rev. 32, e00138–18 (2019).

7. Pang, Z., Raudonis, R., Glick, B. R.Lin, T.-J. & Cheng, Z. Antibiotic resistance in Pseudomonas aeruginosa: mechanisms and alternative therapeutic strategies. Biotechnol. Adv. 37, 177–192 (2019).

8. Langendonk, R. F., Neill, D. R. & Fothergill, J. L. The Building Blocks of Antimicrobial Resistance in Pseudomonas aeruginosa: Implications for Current Resistance-Breaking Therapies. Front. Cell. Infect. Microbiol. 11, 665759 (2021).

9. Crull, M. R. et al. Changing Rates of Chronic Pseudomonas aeruginosa Infections in Cystic Fibrosis: A Population-Based Cohort Study. Clin. Infect. Dis. Off. Publ. Infect. Dis. Soc. Am. 67, 1089–1095 (2018).

10. Feliziani, S. et al. Coexistence and Within-Host Evolution of Diversified Lineages of Hypermutable Pseudomonas aeruginosa in Long-term Cystic Fibrosis Infections. PLOS Genet. 10, e1004651 (2014).

11. Hall, K. M., Pursell, Z. F. & Morici, L. A. The role of the Pseudomonas aeruginosa hypermutator phenotype on the shift from acute to chronic virulence during respiratory infection. Front. Cell. Infect. Microbiol. 12, (2022).

12. Chiang, A. D. et al. Hypermutator strains of Pseudomonas aeruginosa reveal novel pathways of resistance to combinations of cephalosporin antibiotics and beta-lactamase inhibitors. PLOS Biol. 20, e3001878 (2022).

13. Rees, V. E. et al. Characterization of Hypermutator Pseudomonas aeruginosa Isolates from Patients with Cystic Fibrosis in Australia. Antimicrob. Agents Chemother. 63, e02538–18 (2019).

14. Hatfull, G. F., Dedrick, R. M. & Schooley, R. T. Phage Therapy for Antibiotic-Resistant Bacterial Infections. Annu. Rev. Med. 73, 197–211 (2022).

15. Alipour-Khezri, E., Skurnik, M. & Zarrini, G. Pseudomonas aeruginosa Bacteriophages and Their Clinical Applications. Viruses 16, 1051 (2024).

16. Brandão, A. et al. Differential transcription profiling of the phage LUZ19 infection process in different growth media. RNA Biol. 18, 1778–1790 (2021).

17. Zhang, Y., Gao, J., Huang, Y. & Wang, J. Recent Developments in Single-Cell RNA-Seq of Microorganisms. Biophys. J. 115, 173–180 (2018).

18. Gupta, A. et al. Dynamics of phage-host interactions in Bacteroides fragilis resolved by single-cell transcriptomics. Nat. Commun. 17, 4035 (2026).

19. Brennan, M. A. & Rosenthal, A. Z. Single-Cell RNA Sequencing Elucidates the Structure and Organization of Microbial Communities. Front. Microbiol. 12, (2021).

20. Wang, B. et al. Single-cell massively-parallel multiplexed microbial sequencing (M3-seq) identifies rare bacterial populations and profiles phage infection.Nat. Microbiol. 8, 1846–1862 (2023).

21. Blattman, S. B., Jiang, W., Oikonomou, P. & Tavazoie, S. Prokaryotic single-cell RNA sequencing by in situ combinatorial indexing. Nat. Microbiol. 5, 1192–1201 (2020).

22. Ma, P. et al. Bacterial droplet-based single-cell RNA-seq reveals antibiotic-associated heterogeneous cellular states. Cell 186, 877–891.e14 (2023).

23. Kuchina, A. et al. Microbial single-cell RNA sequencing by split-pool barcoding. Science 371, eaba5257 (2021).

24. Samanta, P., Cooke, S. F., McNulty, R., Hormoz, S. & Rosenthal, A. ProBac-seq, a bacterial single-cell RNA sequencing methodology using droplet microfluidics and large oligonucleotide probe sets. Nat. Protoc. 19, 2939–2966 (2024).

25. Ceyssens, P.-J. et al. The Phage-Encoded N-Acetyltransferase Rac Mediates Inactivation of Pseudomonas aeruginosa Transcription by Cleavage of the RNA Polymerase Alpha Subunit. Viruses 12, 976 (2020).

